# State dependent motor cortex stimulation reveals distinct mechanisms for corticospinal excitability and cortical responses

**DOI:** 10.1101/2024.07.15.603619

**Authors:** Nipun D Perera, Miles Wischnewski, Ivan Alekseichuk, Sina Shirinpour, Alexander Opitz

**Affiliations:** Department of Biomedical Engineering, University of Minnesota, Minneapolis, MN, USA

## Abstract

Transcranial magnetic stimulation (TMS) is a non-invasive brain stimulation method which can modulate brain activity by inducing electric fields in the brain. It is a popular tool to study causal brain-behavior relationships. However, brain states vary over time and affect the response of TMS. Neural oscillations can track the current brain state and are a promising marker to guide stimulation timing. Real-time, state-dependent brain stimulation has shown that neural oscillation phase modulates corticospinal excitability reflecting the connection from the primary motor cortex to a target muscle. However, such motor-evoked potentials (MEPs) only indirectly reflect motor cortex activation and are unavailable at other brain regions of interest. The direct and secondary cortical effects of phase-dependent brain stimulation remain an open question. In this study, we recorded the cortical responses during single-pulse transcranial magnetic stimulation (TMS) using electroencephalography (EEG) concurrently with the MEP measurements. TMS was delivered at peak, rising, trough, and falling phases of mu (8-13 Hz) and beta (14-30 Hz) oscillations in the motor cortex. The cortical responses were quantified through TMS-evoked potential components N15, P50, and N100 as peak-to-peak amplitudes (P50-N15 and P50-N100). We further analyzed whether the pre-stimulus frequency band power was predictive of the motor cortical responses. We found a significant main effect of neural oscillation phase on early evoked component (P50-N15). Furthermore, we found an interaction effect of oscillation phase and frequency on both early and late (P50-N100) components. Next, we compared the direct EEG response to the corticospinal excitability reflected by MEP amplitude. Interestingly, the preferred phase of the mu rhythm showed a 90^0^ phase shift between the early TEP components and MEPs. The late component showed the same phase preference between EEG and MEPs. However, such a well-defined relationship did not exist for either of the components during beta phase specific stimulation. In addition, pre-TMS mu oscillatory power and phase significantly predicted both early and late cortical EEG responses when mu rhythm was targeted, indicating the independent causal effects of phase and power. However, only pre-TMS beta power significantly predicted the early and late TEP components when beta rhythm was targeted. Further analysis indicated that both pre-TMS mu and beta power jointly affect early cortical responses. In contrast, the late cortical responses were only influenced by pre-TMS mu power. These findings provide insight to mechanistic understanding of neural oscillation states in cortical and corticospinal activation in humans.

## 1. Introduction

Transcranial magnetic stimulation is a noninvasive brain stimulation technique that can safely modulate brain activity in humans. TMS is widely used in research as a perturbation tool to study causal brain-behavior relationships. TMS induces electric fields in the targeted region of the brain, which results in local modulation of neural activity and propagated downstream effects. As the brain is dynamic in nature, meaning that neural, regional, and metabolic activity is changing in each moment, TMS effects depend on the current brain state. The perturbation of brain with TMS at different states allows us to explore the relationship between brain state and physiological response. Neural oscillations measured from scalp using electroencephalography can reflect various brain states and can be categorized into several frequency bands of interest [1–3]. These oscillations result from neural membrane de-and hyper-polarization and are thus related to local brain excitability [4–6]. Therefore, the phase of a neural oscillation is an indicator of the brain state [7–10]. External stimulation using TMS at different oscillation phases will thus occur at variable levels of brain excitability. However, the exact relationship between oscillation phases and brain responses to TMS is not fully understood.

Recent advances have allowed for testing the causality between oscillation phase of the mu (8-13 Hz) and beta (14-30 Hz) rhythm and cortical excitability in the primary motor cortex (M1). TMS can be applied on M1 to contract a target muscle via corticospinal activation. Through real-time readout and analysis of EEG, the application of TMS can be executed at a particular oscillation phase. Studies using real-time TMS-EEG have demonstrated and replicated that M1-muscle corticospinal excitability depends on oscillation phase [16–19]. While the trough and rising phase of the mu rhythm results in the largest activation by TMS, for the beta rhythm the peak and falling phase were related to the largest MEP response [16,19–22].

While previous findings have shed light on the relation between oscillations and brain excitability, our understanding remains incomplete. Specifically, corticospinal excitability is measured by investigating a TMS-induced muscle response, referred to as a motor-evoked potential (MEP). As such, this method is restricted to the motor cortex and is unable to study phase-dependencies in other parts of the cortex. Furthermore, while they are indicative of cortical activation, MEPs are confounded by background activity fluctuations at the spinal and muscle level. To overcome these limitations, the phase-specific effects of TMS can be measured on EEG potentials (referred to as TMS-evoked potentials [TEPs]) more directly. In addition, TEPs can be recorded from other locations on the scalp and can thus be used for non-motor regions as well. TEPs reflect a broad range of activity in the targeted brain region. Early TEPs in the motor cortex, for example, N15, reflect direct cortical activation of the motor/premotor area [23–25]. The later components such as P50 and N100 may reflect reafferent activity and secondary effects from multiple sources [22,25–27]. Previous non-human primate studies by our group have shown these direct and peripherally induced TMS effects can be dissociated using invasive electrophysiology [28] highlighting their potential to measure direct cortical responses to TMS. Currently, there is sparse evidence for the effects of oscillation phase on TEPs. One study showed a phase-dependent modulation of early and late TEP components (e.g. P60/P70 and N100) during peak and trough phases of the mu rhythm [22]. TEPs in the beta rhythm have not been investigated so far to our knowledge.

The goal of the present study is 1) to systematically investigate the causal relationship between cortical activation, as measured by TEPs, and oscillation phase in the mu and beta rhythm; 2) to directly compare the effects of real-time phase-dependent TMS on cortical activation, as measured by TEPs, and corticospinal excitability, as measured by MEPs; and 3) study the role of pre-TMS power in TEP generation. Our results indicate phase-dependent modulation of early and late TEP components in both mu and beta oscillations. Furthermore, corticospinal excitability and cortical activity follow a 90^0^ out of phase relationship for mu oscillation while beta did not show a quantifiable relationship. We also find that early TEP components are modulated by sensorimotor mu and beta rhythms whereas the late component is modulated by mu rhythm alone. Our results indicate the possibility of distinct mechanisms that modulate corticospinal excitability and cortical responses in the sensorimotor cortex. This is important when developing biomarkers of brain function and excitability, an important consideration in clinical outcomes at large.

## 2. Methods

### 2.1. Participants

We recruited 20 healthy volunteers for this study (11 female, mean ± std age: 22.7 y ± 2.9, range: 18-45 y). All participants were right-handed and without reported history of neurological or psychiatric disorders. The study was approved by the institutional review board of the University of Minnesota and all participants were consented prior to participation in the study. The reader is referred to Wischnewski et al. for additional information on the dataset used in this study [29].

### 2.2. Transcranial Magnetic Stimulation and real-time triggering

We used Magstim Rapid^2^ stimulator with a figure-of-eight D70^2^ coil (Magstim Inc., Plymouth, MN, USA) to deliver single pulse TMS. We recorded electromyography (EMG) from the right first dorsal interosseus (FDI) muscle using BIOPAC ERS100C amplifier (BIOPAC systems Inc., Goleta, CA, USA). Initially, the coil was oriented at 45^0^ relative to the midline and the hotspot was determined as the coil location and orientation eliciting the highest motor evoked potential (MEP). Once the hotspot was located, the coil location and orientation were continuously tracked using Brainsight neuronavigation system (Rogue Research, Montreal, Canada). The resting motor threshold (rMT) was determined based on maximum likelihood parameter estimation by sequential testing (PEST) [30]. For the subsequent real-time TMS, the intensity was set to 120% rMT.

### 2.3. EEG acquisition and real-time TMS triggering

We recorded the EEG using 64-channel actiCAP Slim active electrodes and actiCHamp EEG amplifier (Brain Products GmbH, Gilching, Germany). For this study, we used the educated temporal prediction (ETP) algorithm [31] to stimulate rising, peak, falling, and trough phases of sensorimotor mu (8-13 Hz) and beta (14-30 Hz) oscillations. The ongoing oscillation for phase targeting was calculated as the Laplacian C3 signal with 8 surrounding electrodes (FC1, FC3, FC5, C1, C5, CP1, CP3, and CP5). Resting state EEG was acquired for 3 minutes immediately prior to the stimulation to train the ETP algorithm. Subsequently, the same Laplacian C3 signal was used for real-time TMS triggering, while ongoing EEG activity was recorded and saved at the end of each experiment session.

Each participant underwent two separate experiment sessions for targeting mu and beta oscillations. The sessions were at least 48 hours apart to prevent carryover effects and were randomized across participants. During each experiment session, the participant received a total of 600 pulses, in four blocks of 150 pulses. In each block, the phases were targeted pseudorandomly, where both the experimenter and the participant were blinded to the target phase. The pulses were delivered with a jitter of 2-3 s to prevent the effects of last pulse on the current pulse.

### 2.4. Data processing and statistical analysis

#### 2.4.1. Electrophysiological data processing

The MEPs were extracted from a window of 20-60 ms following TMS pulse. MEPs were considered noisy if the average absolute baseline EMG activity (−100 ms to 0 ms relative to TMS pulse) was above 0.02 mV and larger than absolute average EMG activity + 2.5 times standard deviation of resting state EMG. sufficiently deviated from a predefined threshold. For each subject, the individual MEPs were normalized to the average of all MEPs.

EEG data was preprocessed using custom scripts in MATLAB (MathWorks Inc., Natick, MA, USA) using FieldTrip toolbox [32]. The raw EEG data from 4 blocks was combined into one block of 600 trials. This EEG data was separated into epochs between -1 s and 1 s, time-locked to TMS trigger (t = 0). The data was then demeaned and detrended, and the pulse artifact and early muscle activity between -6 ms and 10 ms was removed by padding the time window. The excluded time window was then interpolated using the cubic Hermite interpolating polynomial (pchip). The bad channels and trials were then removed by visual inspection. On average, 6.6% of the trials for targeting mu and 8.0% of the trials for targeting beta were removed (7.3% trials removed from the total number of trials). On average, one channel was removed from each session for each participant. We performed independent component analysis (ICA) to remove residual muscle activity, and ocular artifacts using the infomax ICA algorithm [33] implemented in FieldTrip. On average, 6.7 components were removed from mu session and 6 components were removed from beta session. The data was then resampled to 1 kHz and bandpass filtered between 1 and 50 Hz using a 3^rd^ order Butterworth filter. We interpolated the channels using spline interpolation of neighboring channels. The definition of neighboring channels was acquired from the default template available in FieldTrip for the EEG cap used in the experiment. Finally, the data was referenced to the common average and baseline corrected with a window of -500 ms to -15 ms.

Once EEG data was preprocessed, we clustered the trials based on the targeted phase and calculated the average TMS evoked potentials by averaging across trials. We identified the TEP components that are commonly reported in literature, N15, P30, P60, and N100 [34]. While N15 and N100 were consistent, the prominence of P30 and P60 peaks varied across participants. Some participants showed a pronounced P30-N45-P60 complex, while the others showed either P30 or P60. For our analysis, we extracted the peak TEP components from predefined time windows of interest, 10–20, 20–70 and 90– 130 ms. In the 20–70 ms window, we extracted the most prominent peak in each participant. We called these M1 TEP peaks, N15, P50 (despite the variation of latency in the 20–70 ms time window), and N100. Given that these evoked potentials are noisy in the trial level, we defined N15 potential as the mean amplitude 15 ± 2 ms after the TMS pulse, P50 potential – 50 ± 5 ms, and N100 potential – 100 ± 10 ms for the trial level analysis of TEP components. These windows were chosen considering the prominence of the peaks.

Pre-TMS power spectrum of C3 electrode was calculated to investigate whether pre-stimulus activity correlates to post-stimulus evoked activity. We did so by applying fast Fourier transform with a single Hanning taper at a resolution of 1 Hz on the time window between -550 and -50 ms. Pre-TMS mu and beta power were calculated as the mean power in the 8-13 and 14-30 Hz range respectively.

#### 2.4.2. Interpretability of TEPs in a phase-locked regime

The interpretation of TEP components when TMS is delivered at a specific phase repetitively, comes with a limitation. While the time-locked averaging procedure cancels out background EEG activity, phase-locked activity survives. The amplitudes of TEP components could be interpreted only if TMS causes transient phase resetting of the ongoing oscillation. If such a transient phase reset does not occur, a correction could be applied by creating phase-locked non-stimulated trial and channel-wise subtraction of phase-locked non-stimulated average from phase-locked stimulated average [22,35,36]. We tested whether a transient phase reset occurs during TMS using the measure, phase preservation index (PPI) [22,37]. We performed this by offline targeting of phases on the resting data to create non-stimulated trials in 18 sessions out of 40 (9 per mu and beta). The resting data for the other 22 sessions was not available due to an error related to saving data. However, we did not find conclusive evidence for stability of phase following TMS delivery (Supplemental Figure S1). Since we cannot conclusively prove that phase is reset during TMS, we utilized peak-to-peak amplitudes of TEP components to quantify the cortical responses. Therefore, we considered two TEP complexes in our study, P50-N15 (early response) and P50-N100 (late response).

#### 2.4.3. Statistical analysis

In group level analysis of the TEP amplitudes, repeated-measures analysis of variance (rmANOVA) was performed on target phase and rhythm. This was followed by multiple comparisons tests to investigate pairwise differences according to Tukey HSD correction.

To investigate the relationship between TEP and MEP relationship, we used a preferred phase approach. We did so by reducing the electrophysiological measures to two orthogonal components (peak-trough and fall-rise). We then calculated the resultant magnitude and direction by converting the two orthogonal components into polar coordinates. To test whether MEP and TEP components have a predictable phase dependency, we calculated the phase difference between the preferred phase of MEP and TEP components. A phase difference of 0^0^ would result in a similar phase relationship, whereas a phase difference of 90^0^ would mean an opposite relationship. We calculated this between MEP and each TEP component for each participant.

To study the effect of pre-TMS band power on TEP components, we ran generalized linear mixed-effect models on trial level each target band. We considered the target phase (rising, peak, falling, trough) and target pre-TMS band power as fixed effects, and participant number as random effect. On the group level, we studied the relationship between pre-TMS power and average TEP amplitude by calculating Spearman’s rank correlation.

We further investigated how the interaction of pre-TMS mu and beta power affect the cortical responses. We pooled TEP and power data from all trials (from both mu and beta target sessions) and performed a median split on the mu and beta power separately for each participant. Next, we classified the trials into four classes as mu high-beta high, mu high-beta low, mu low-beta high, mu low-beta low. Finally, we calculated the class average of early and late responses for each participant and normalized it to the subject average. We performed Wilcoxon signed rank test to compare the classes in a pairwise manner. An additional analysis was performed on pre-TMS delta and theta rhythms separately using the same approach. This was done to verify that TEP responses are not modulated by the pre-TMS delta and theta power. For all analyses, the significance level was set to be α = 0.05.

## 3. Results

### 3.1. Phase dependent modulation of TEP amplitude

First, we quantified the early (P50-N15) and late (P50-N100) peak-to-peak TEP complexes in the time-locked averaged signal per participant. TMS delivered at specific phases of oscillations shows differential modulation of cortical responses, dependent on target phase and oscillatory frequency. We observed that mu-specific targeting resulted in larger amplitude changes in early (0.95 – 1.06 n.u.) and late (0.97 – 1.03 n.u.) cortical responses. Comparatively, beta-specific targeting resulted in lesser changes in early (0.98 – 1.02 n.u.) and late (0.99 – 1.01 n.u.) cortical responses (Fig. 2). This suggested generally stronger effects at the mu rhythm. Similarly, the early response showed larger phase dependent modulations compared to the late response for both target rhythms.

**Figure 1:**
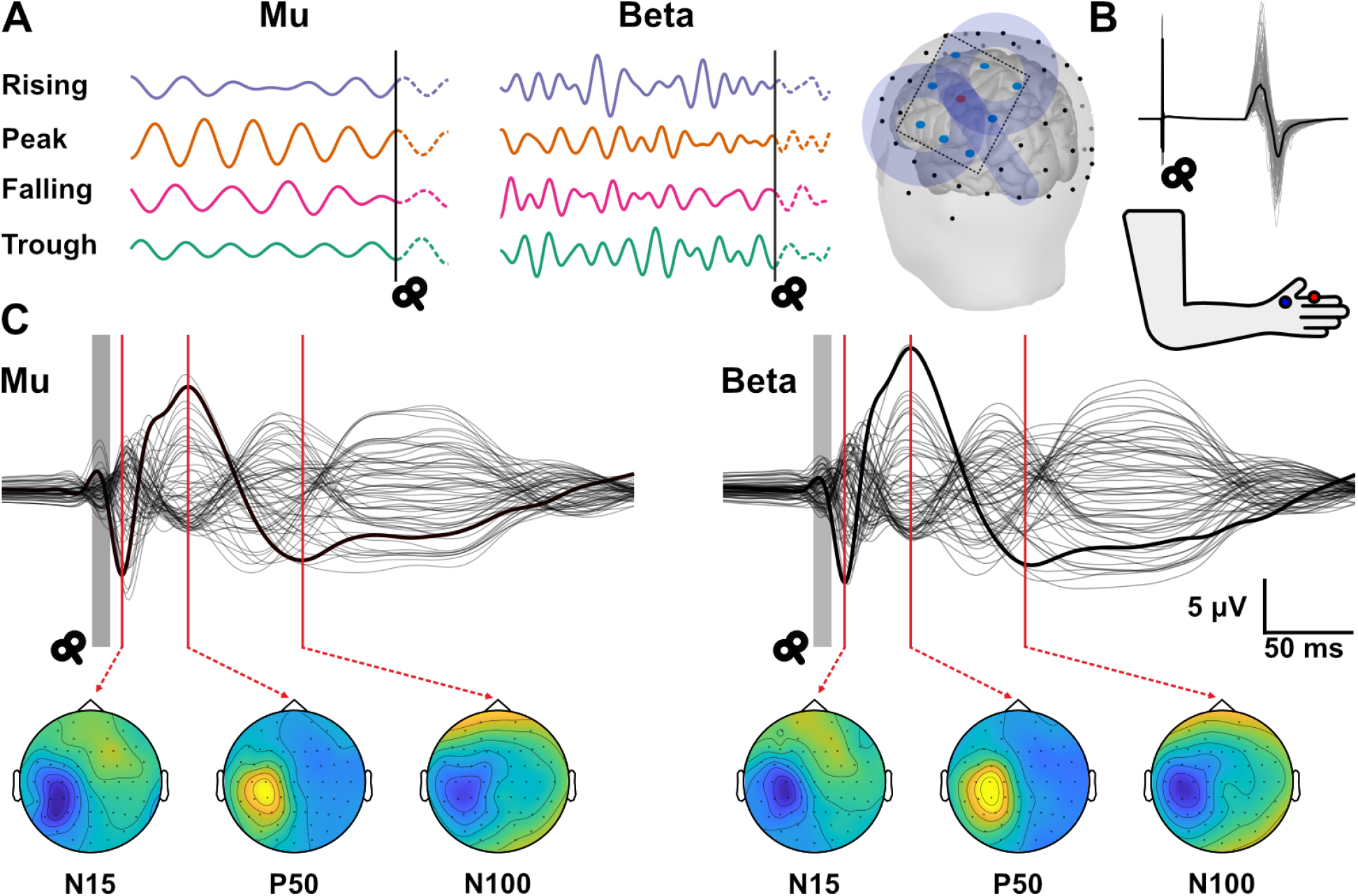
The overview of phase and frequency targeting, and TMS evoked potentials (TEPs). A) An example of phase targeting during the training phase of the educated temporal prediction (ETP) algorithm. ETP algorithm accurately targets rising, peak, trough and falling phases of both mu and beta frequency bands. The rendering of the brain shows the Laplacian montage of the electrodes (FC5, FC3, FC1, C5, C1, CP5, CP3, and CP1) used to extract the C3 signal. The coil was placed on the motor hotspot corresponding to first dorsal interosseus (FDI) muscle. B) Example of motor evoked potentials (MEPs) acquired from FDI muscle for a single participant with bold trace representing the mean MEP. Bottom panel shows the montage used for EMG acquisition from FDI muscle. C) The global average TEP signal for each target frequency band. The bold trace indicates the C3 signal. The topography of the three components, N15, P50 and N100 used to characterize cortical responses are shown below each trace.

**Figure 2:**
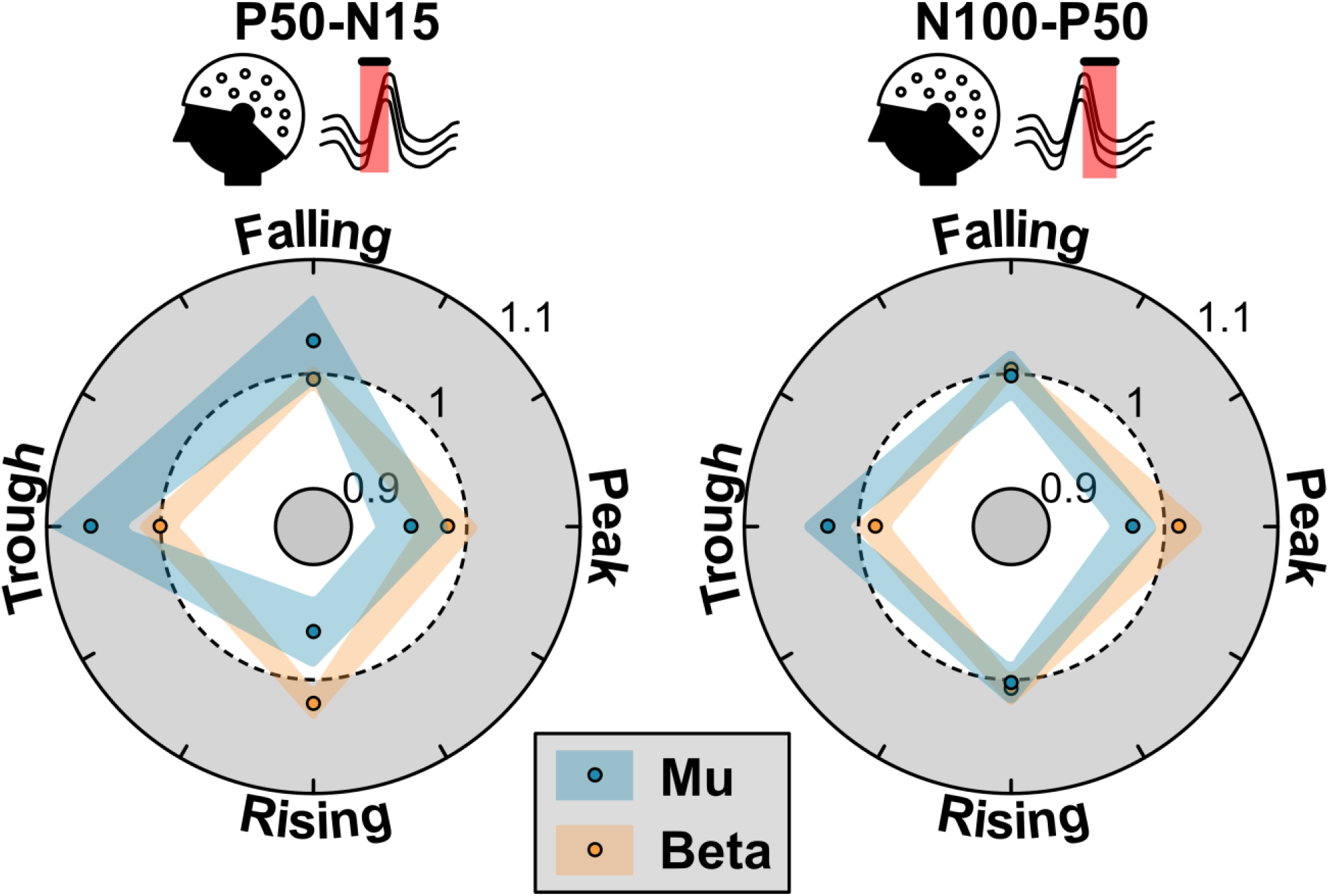
Phase dependency of TEP components for mu and beta rhythms. Left: Phase dependency of the early component, P50-N15. Mu target rhythm showed significant increased modulation of TEP amplitude for trough and falling phases compared to peak and rising (p < 0.02). For beta target rhythm, rising phase showed significantly higher TEP amplitude modulation compared to peak phase (p = 0.02). Right: Phase dependency of the late component, P50-N100. The late component showed opposite amplitude modulation for mu and beta phases where mu showed significant increase of amplitude for trough phase compared to peak phase (p = 0.001) and vice versa for mu phase (p = 0.04). The shaded area represents the 95% confidence interval.

Next, we investigated the effects of target phase and target rhythm on the early TEP component. At the group level, rmANOVA showed a significant main effect of target phase [F(3, 57) = 6.96, p < 0.001] and a significant target phase × target oscillation interaction (F(3, 57) = 5.96, p = 0.001) on the early TEP component. When mu-specific target is considered, trough to falling phases showed increased modulation (1.03 – 1.06 n.u.) of cortical early response. Conversely, peak and rising phases showed decreased modulation (0.95 – 0.96 n.u.). The difference between increased and decreased modulation was statistically significant (post-hoc multiple comparisons test with Tukey HSD correction; p < 0.02). During beta-specific targeting, the early response showed increased modulation at rising phase (1.02 n.u.) and decreased modulation at peak phase (0.98 n.u.). The difference between these phasic modulations was significant (p = 0.02).

We then investigated the effects of target phase and target rhythm on the late response. The late response did not show a significant main effect of phase; however, it showed a significant target phase × target oscillation interaction [F(3, 57) = 5.66, p = 0.013]. The late component showed an opposing phase dependent modulation for mu and beta rhythms. For mu-specific target, trough phase resulted in the increased modulation of the late response (1.03 n.u.) while peak phase resulted in the decreased modulation (0.97 n.u.). This modulation of the late response between the peak and trough phases was significantly different (p = 0.001). During beta-specific targeting, peak phase resulted in the increased response modulation (1.01 n.u.) while trough phase resulted in the decreased response modulation (Fig. 2B). Likewise, for beta-specific target, the modulation between peak and trough phases was statistically significant according to multiple comparisons (p = 0.04).

Overall, this showed higher phasic modulation of both early and late responses for mu-specific targeting compared to beta-specific targeting. Similarly, early responses showed larger changes compared to late responses. This underscores the roles of mu and beta phase on TMS evoked cortical responses at different latencies.

### 3.2. MEP and TEP phase relationship

Previously, we found that MEP excitability for mu and beta target oscillations have an opposing phase relationship at the group level. Wischnewski et al., 2022 showed mu-specific targeting results in high excitability for trough and rising phases. Conversely, beta-specific targeting results in high excitability for peak and fall in beta oscillations. Here, we calculated the preferred phase of the MEPs and TEP components for each participant to quantify the phase - excitability relationship. As expected, the distribution of the preferred phase was oriented towards trough and rising phases in mu and peak and fall phases in beta oscillations for MEPs. In the TEP analysis, the distribution of the preferred phase of the early TEP component was oriented towards falling and trough phases. For late TEP component, the orientation was dominant towards trough and rising phases (Fig. 3A and Fig. 4). This indicated that the early TEP component for mu target rhythm showed a 90^0^ shift in the preferred phase relative to MEPs, whereas the late component showed the same phase preference as MEPs. However, neither the early nor late TEP component following beta specific stimulation showed a strong phase preference towards a particular orientation (Fig. 3B and Fig. 4).

**Figure 3:**
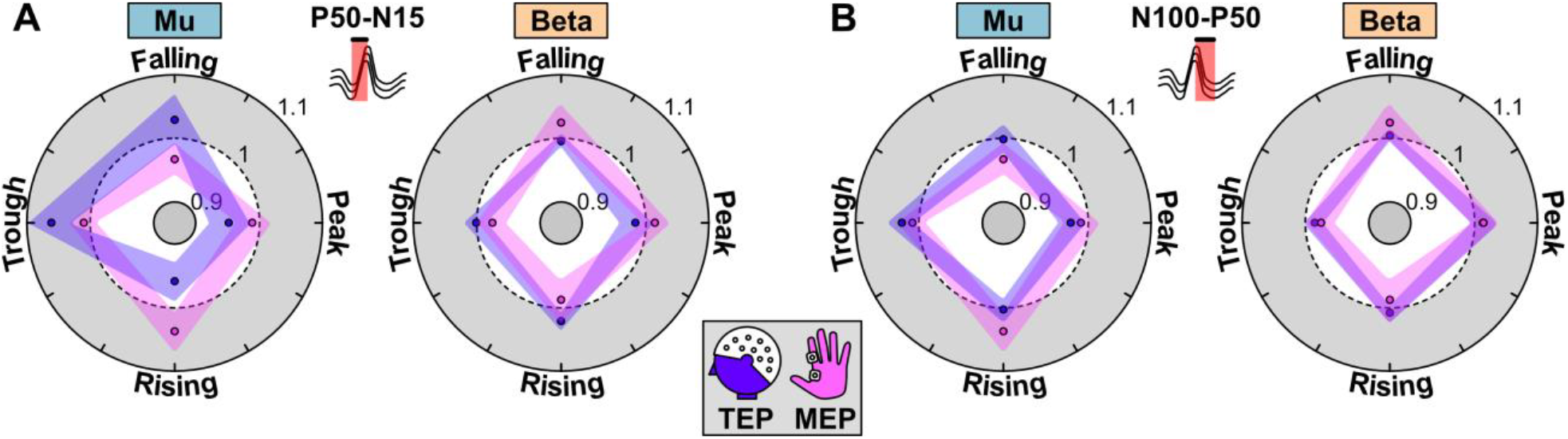
Relationship between corticospinal excitability and cortical responses. A) The comparison of relative changes of MEP and TEP amplitudes for the early component for mu and beta oscillations. B) The same comparison for the late component.

**Figure 4:**
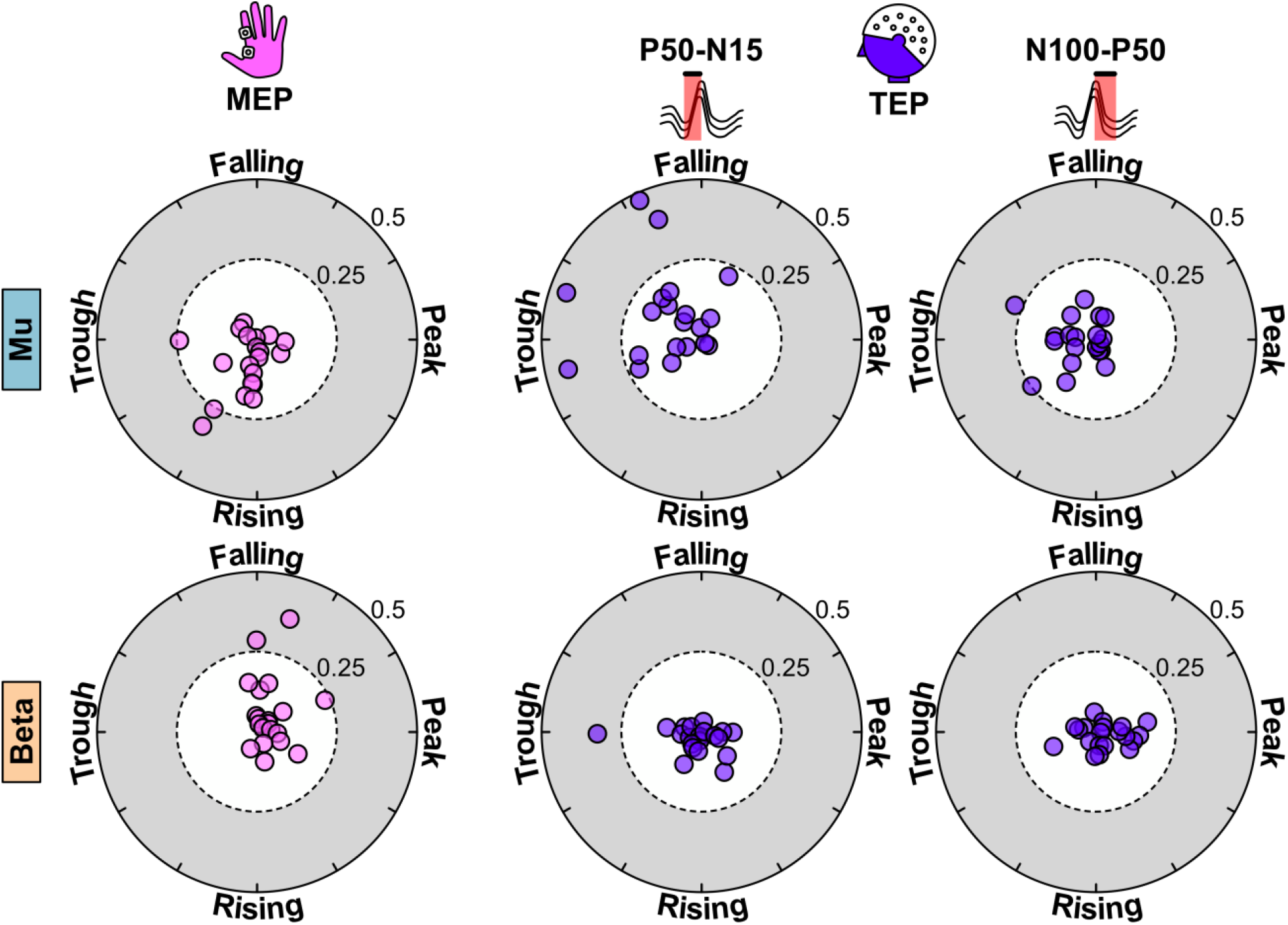
Individual phase preferences of MEP and TEP components. (Top) MEPs generated during mu target rhythm have a phase preference of rising and trough phases while the early TEP component is preferentially modulated by falling and trough phases. The late component shows a phase preference similar to that of MEPs where the preferred orientation is towards trough and rising phases. (Bottom) MEPs generated when phases of beta rhythm were targeted, are preferentially modulated by falling and peak phases (opposite to mu). Neither the early component nor the late component showed a preferential modulation of TEP amplitude to a specific phase.

### 3.3. Effect of pre-TMS power on cortical responses

We ran two GLMMs for each target rhythm to study the effects of target phase and the pre-TMS band power on early and late components. For mu rhythm, we found significant main effects of phase (early component: F = 69.37, p < 0.001; late component: F = 21.43, p < 0.001) and pre-TMS mu power (early component: F = 92.98, p < 0.001; late component: F = 25.54, p < 0.001) and an interaction effect of phase × pre-TMS power (early component: F = 41.00, p < 0.001; late component: F = 32.18, p < 0.001) for both early and late TEP components. Interestingly, we only found a significant effect of pre-TMS beta power, but not phase, on the early and late TEP components when beta rhythm was targeted (early component: F = 9.66, p = 0.002; late component: F = 11.77, p < 0.001).

To study whether pre-TMS power is related to MEP amplitude and TEP amplitudes at the group level, we calculated the correlation between these measures. We found that the early and the late components during mu target rhythm were positively correlated with the pre-TMS power (early component: r = 0.57, p = 0.008 and late component: r = 0.76, p < 0.001) (Fig. 5A). For beta target rhythm, the early component showed a significant positive correlation with the pre-TMS beta power, but not the late component (early component: r = 0.45, p = 0.048 and late component: r = 0.42, p = 0.067) (Fig. 5). The strength of correlations suggests that pre-TMS mu power is the stronger determinant of the brain responses between the two dominant oscillations in the sensorimotor cortex.

**Figure 5:**
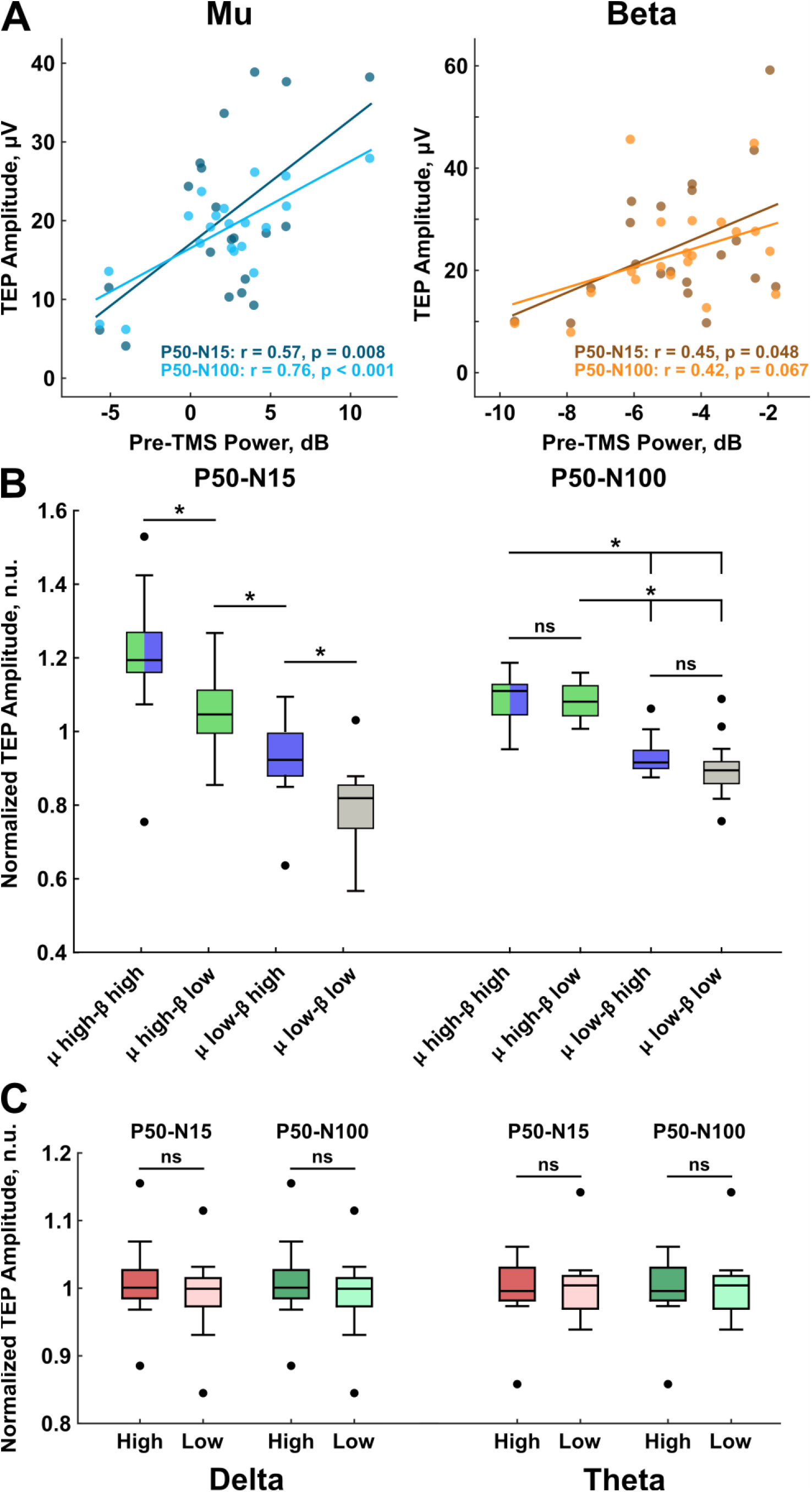
The effects of pre-TMS power on TEP components. A) The group level correlation between pre-TMS power and TEP components. Pre-TMS mu (8-13 Hz) band power showed a strong correlation with both early (r = 0.57, p = 0.008) and late (r = 0.76, p < 0.001) TEP components (left). Pre-TMS beta (14-30 Hz) bad power showed a moderate correlation with the early TEP component (r = 0.45, p = 0.048); however, the correlation was not significant for the late component (r = 0.42, p = 0.067). Each dot in the scatter plot represents the average power and average TEP amplitude across the four phases per participant. B) The combined effect of pre-TMS mu and beta power on the TEP components. The early component showed highest modulation when both mu and beta power were high and lowest modulation when both mu and beta power were low. High mu-low beta and low mu-high beta resulted in higher than average and lower than average modulation respectively. Wilcoxon rank sum tests showed that the relative changes of TEP amplitudes in these groups were significantly different (left). The modulation of late component was not affected by beta power. Regardless of beta power, high mu power resulted in higher TEP modulation and low mu power resulted in lower TEP modulation. Significant differences were not observed within groups with high mu power or low mu power. However, differences were significant across groups with high mu power and low mu power (right). C) The power of other bands of interest, delta and theta, did not contribute to modulation of both early and late components. Outliers are marked in black circles. * p < 0.05, n.s. = not significant.

We further studied the combined effects of pre-TMS mu and beta power on the early and late components by pooling the trials from all sessions and classifying the mu and beta power as ‘high’ and ‘low’ via a median split. Hence, the TEPs were divided into four classes: mu high - beta high, mu high - beta low, mu low - beta high, mu low - beta low. The comparisons across groups indicate that both mu and beta power affected the early TEP component with increasing power resulting in increased TEP modulation (Fig. 5B). We performed Wilcoxon rank sum test to compare classes and found that they were pairwise significant (mu high-beta high vs. mu high-beta low: z = 2.90, p = 0.004, mu low-beta high vs. mu low-beta low: z = 3.70, p < 0.001). Interestingly, only the mu power and not the beta power showed an effect on the late component (Fig. 5B). The difference of late TEP modulation between mu high-beta high and mu high-beta low classes were not significant (z = 0.60, p = 0.55). Similarly, mu low-beta high and mu low-beta low groups did not show a significant difference (z = 1.90, p = 0.06). However, both classes with high mu power showed significantly higher modulation of late TEP component compared to both classes with low mu power irrespective of the beta power (mu high-beta high vs. mu low-beta high: z = 3.70, p < 0.001, mu high-beta high vs. mu low-beta low: z = 3.58, p < 0.001 and mu high-beta low vs. mu low-beta high: z = 3.78, p < 0.001, mu high-beta low vs. mu low-beta low: z = 3.66, p < 0.001). In order to verify that the power of other frequency bands (delta and theta) do not contribute to TEP modulation, we classified TEPs into high and low power states of these bands. However, the results did not indicate significant modulation of TEP amplitudes for early or late components in both delta and theta bands. Hence, the results indicate that early response is driven by a combination of mu and beta power whereas the late component is driven by mu power alone.

## 4. Discussion

In this study, we investigated the effects of state-dependent stimulation of the motor cortex on cortical responses for mu and beta oscillations and their relationship to corticospinal excitability. We defined an immediate oscillatory state as phase and power of oscillation of interest. The oscillatory phase was extracted in real-time for ad hoc stimulation, while pre-TMS band power was estimated in a post-hoc manner. First, we found that the early cortical responses (<50 ms) were modulated by mu and beta phase during the TMS pulse. The later component (50-100 ms) also showed modulation of cortical responses, albeit to a lesser degree. Overall, mu-specific targeting elicited larger changes in amplitudes compared to beta. Second, we found that the M1-muscle excitability measured through MEPs showed a different phase preference to that of the early cortical response in mu-specific targeting. For the same target rhythm, the phase preference between later cortical activation and MEPs was similar. Finally, we found that pre-TMS mu and beta power affect early cortical responses independent of each other. However, only pre-TMS mu power affects the later cortical response.

Previous studies have found that N15 is indicative of early excitation of motor and premotor areas [23–25], P50 reflects the reafferent signals from the stimulated muscle (motor-basal ganglia cortical loop) [22,25,26], and N100 is related to GABAergic inhibition and cortical silent period [38–43]. In addition to these components, P30 and N45 are reported [44,45]. However, individual variability exists pertaining to these components. In our study, several participants revealed the P30-N45-P60 complex (we denote the second positive peak in this complex as P50 considering the group level latency). This individual variability meant that it was not a reliable measure at the group level. Consequently, the global response did not delineate P30 and N45 components, thus we constrained our analysis to P50 component. Our study shows that the strength of early cortical excitation and muscle reafferent signal, as estimated by P50-N15 component, is highly modulated by the phase of the mu oscillation. We observed that trough and falling phases of the mu oscillation caused increased modulation of this response. Targeting beta oscillation modulated this to a lesser degree, with rising phase showing increased activity. However, the physiological processes associated with the late components showed opposing effects for mu and beta rhythms. We also found that the early component was more affected by phase targeting compared to the late component. This implies that the phase affected early activation which correlates to MEP generation more than the later components. The strong modulation of early component by mu phase thus indicates that mu rhythm is a driving factor of local brain excitability. The contribution of beta rhythm to this modulation is weaker according to our results. The observation that the overall relative changes of the TEP components were greater for mu compared to beta oscillations could also corroborate the importance of mu rhythm in cortical response generation.

A key finding in this study is that the modulation of early cortical response (as measured by TEPs) does not resemble the same pattern as the M1-muscle excitability for mu-specific target (as measured by MEPs). Instead, the early direct cortical activation and the muscle excitability show a different phase preference that is 90^0^ out of phase to each other. This could be explained by the cortical origins of early TEP components and MEPs. Sensor-level and source level analysis of TEP components have shown widespread activity in M1, supplementary motor area, and superior parietal lobule during the early response [24,46]. On the other hand, MEPs from FDI muscle have been shown to be generated by gyral crown and upper parts of the sulcal wall in M1 [47,48]. Essentially, early response captured by C3 represents the aggregate activity from a wider cortical area. In contrast, the MEPs represent the activity of a small subset of neurons that have a specific orientation. Hence, we could expect this subset of neurons to be modulated by a phase of ongoing oscillation that is different from overall population activated by TMS. Interestingly, the preferred phase of both later cortical response and M1-muscle excitability was oriented towards trough and rising phases in mu-specific targeting. As previously discussed, the late components reflect the reafferent information from the motor-basal ganglia-cortical loop (P50) and cortical inhibition and cortical silent period (N100). Given that trough and rising phases indicate a high M1-excitability state, the feedback mechanisms may lead to increased inhibition [27,49]. This would result in increased late response for the same phases. Therefore, the early response is likely related to TMS pulse itself, whereas the late components are an after effect of MEP generation. We did not find an alignment of the preferred phases for beta rhythm. This suggested an influence of beta phase on corticospinal excitability and cortical responses that was difficult to reconcile.

We further analyzed whether pre-TMS band power explains the variability of the TEP amplitudes at the trial and the group level. At trial level, both pre-TMS mu power and the target phase of mu oscillation affected brain responses. However, when targeting beta oscillations, only pre-TMS beta power had an effect on brain responses. In our previous study, Wischnewski et al. showed that both phase and pre-TMS mu power (when mu oscillations were targeted) play a role in corticospinal excitability as measured by MEPs. For beta-specific targeting, only phase determined corticospinal excitability. Thus, there is general agreement of the effect pre-TMS mu power and phase on MEPs and TEPs. At the group level, the relationship between pre-TMS band power and TEP components showed a positive association for mu rhythm, confirming the trial level observation. For beta, the early component showed a significant positive association with pre-TMS beta power, but not the later component. We sought to explain the combined effect of pre-TMS mu and beta power on all TEP responses by classifying them into high and low power states. Accordingly, the post-hoc analysis we performed showed that early component is modulated by the power of both mu and beta oscillations. However, only pre-TMS mu power significantly modulated the late component, shedding light on distinct roles of mu and beta in cortical response generation. This confirms the predominant role played by sensorimotor mu oscillatory characteristics in early local excitability and late cortical responses. However, the role of sensorimotor beta characteristics, especially in later cortical responses is unclear. It has been shown that sensorimotor mu oscillations are related to GABA inhibitory activity [16] and inversely related to blood oxygen level dependent (BOLD) signal in the cortical-subcortical motor network [50]. This could explain the positive association between the late component, which is a marker of cortical inhibition and pre-TMS mu power. The effect of both phase and power on corticospinal excitability for sensorimotor mu oscillations has been explored previously for peak and trough phases, which supports the main and interaction effects revealed by the GLMMs [51]. In contrast, the role of sensorimotor beta in the motor cortex is not fully understood. Studies have shown that beta power is associated with voluntary and task related motor activity [52,53] and that resting beta power does not affect the propagation of TMS excitations in the motor cortex [50]. In fact, movement related beta decrease has been shown during spontaneous and trigger movement [52] whereas inhibition of movement leads to higher beta power [53–55]. The fact that the participants in our study did not perform voluntary muscle contractions or motor tasks could explain weaker correlation between the cortical responses and pre-TMS beta power compared to pre-TMS mu power.

It is not fully established if TMS-evoked cortical responses are generated by oscillatory phase-resetting or by adding to the ongoing oscillation. Thus, one could consider if our brain state-dependent TEP results are confounded by pre-TMS voltage differences. Some studies have suggested phase-resetting of ongoing oscillations by TMS or other forms of stimulus presentation [56–59]. However, other studies have found evidence against this that phase is preserved following TMS up to 200 ms through measures of phase preservation index (PPI) [22,37]. To clarify this question, we first calculated the phase preservation index for C3 and surrounding electrodes for real TMS trials and non-stimulated trials, but we were not able to find evidence for phase preservation (Supplementary Fig. 1). Thus, we measured early and later components as a difference between two consecutive TEP peaks (P50-N15 and N100-P50) [22,35,36]. The use of peak-to-peak values mitigates possible confounds of phase preservation or phase resetting.

Here, we only used phase of oscillation as an experimentally controlled variable, while pre-TMS power was estimated post hoc. Studies have shown the importance of power of oscillation to elicit high and low excitability states [51]. In the future, experimental studies may consider delivering TMS based on the combined pre-defined phase and power definitions. It should be noted that we removed the first 15 ms after TMS pulse from the analysis due to TMS pulse artifact. This is a standard approach; however, very early evoked activity around 1-2 ms after TMS delivery is thus not captured by our analysis. Specific experimental settings showed that very early responses may represent activation of pyramidal tract neurons and corticospinal neurons directly and via monosynaptic connections [60].

The present results characterize brain state dependency of motor cortex. An exciting future direction is to extend this work to other brain areas. Furthermore, real-time closed-loop stimulation systems could be developed by taking more state variables, such as band power and network connectivity, into account.

## 5. Conclusion

Here we investigated the effects of brain states quantified by rhythm-specific oscillatory phase and power on TMS evoked activity. We measured the evoked activity through corticospinal excitability (MEP) and cortical responses (TEP). We found that stimulating M1 at specific phases of mu or beta rhythms modulates early and later cortical responses with early responses showing higher modulation. Although both cortical excitability and cortical responses show brain state dependency, the state preferences are different. We also studied the association of pre-TMS power with cortical responses. We found that mu phase and power, and beta power affect TEP components. Further investigation of these power values revealed that both mu and beta power is associated with early TEP responses whereas only mu power is associated with late responses. These findings highlight the importance of real-time, state-dependent brain stimulation, and its relationship to biomarkers. It further makes a compelling case for state-dependent stimulation for pathological conditions to enhance treatment efficacy.

## References

[1] Andrews HL, Jasper HH. Human Brain Rhythms: I. Recording Techniques and Preliminary Results. Journal of General Psychology 1936;15:98.

[2] Dustman RE, Boswell RS, Porter PB. Beta Brain Waves as an Index of Alertness. Science 1962;137:533–4. 10.1126/science.137.3529.533.

[3] Lopes da Silva F. Neural mechanisms underlying brain waves: from neural membranes to networks. Electroencephalography and Clinical Neurophysiology 1991;79:81–93. 10.1016/0013-4694(91)90044-5.

[4] Brienza M, Mecarelli O. Neurophysiological Basis of EEG. In: Mecarelli O, editor. Clinical Electroencephalography, Cham: Springer International Publishing; 2019, p. 9–21. 10.1007/978-3-030-04573-9_2.

[5] Contreras D, Steriade M. Cellular basis of EEG slow rhythms: a study of dynamic corticothalamic relationships. Journal of Neuroscience 1995;15:604–22. 10.1523/JNEUROSCI.15-01-00604.1995.

[6] Schaul N. The fundamental neural mechanisms of electroencephalography. Electroencephalography and Clinical Neurophysiology 1998;106:101–7. 10.1016/S0013-4694(97)00111-9.

[7] Berger B, Minarik T, Liuzzi G, Hummel FC, Sauseng P. EEG Oscillatory Phase-Dependent Markers of Corticospinal Excitability in the Resting Brain. BioMed Research International 2014;2014:936096. 10.1155/2014/936096.

[8] Combrisson E, Perrone-Bertolotti M, Soto JL, Alamian G, Kahane P, Lachaux J-P, et al. From intentions to actions: Neural oscillations encode motor processes through phase, amplitude and phase-amplitude coupling. NeuroImage 2017;147:473–87. 10.1016/j.neuroimage.2016.11.042.

[9] Miller KJ, Hermes D, Honey CJ, Hebb AO, Ramsey NF, Knight RT, et al. Human Motor Cortical Activity Is Selectively Phase-Entrained on Underlying Rhythms. PLOS Computational Biology 2012;8:e1002655. 10.1371/journal.pcbi.1002655.

[10] O’Keeffe AB, Malekmohammadi M, Sparks H, Pouratian N. Synchrony Drives Motor Cortex Beta Bursting, Waveform Dynamics, and Phase-Amplitude Coupling in Parkinson’s Disease. J Neurosci 2020;40:5833–46. 10.1523/JNEUROSCI.1996-19.2020.

[11] Ippolito G, Bertaccini R, Tarasi L, Di Gregorio F, Trajkovic J, Battaglia S, et al. The Role of Alpha Oscillations among the Main Neuropsychiatric Disorders in the Adult and Developing Human Brain: Evidence from the Last 10 Years of Research. Biomedicines 2022;10. 10.3390/biomedicines10123189.

[12] Fingelkurts AA, Fingelkurts AA, Rytsälä H, Suominen K, Isometsä E, Kähkönen S. Composition of brain oscillations in ongoing EEG during major depression disorder. Neuroscience Research 2006;56:133– 44. 10.1016/j.neures.2006.06.006.

[13] O’Reardon JP, Solvason HB, Janicak PG, Sampson S, Isenberg KE, Nahas Z, et al. Efficacy and Safety of Transcranial Magnetic Stimulation in the Acute Treatment of Major Depression: A Multisite Randomized Controlled Trial. Biological Psychiatry 2007;62:1208–16. 10.1016/j.biopsych.2007.01.018.

[14] George MS, Lisanby SH, Avery D, McDonald WM, Durkalski V, Pavlicova M, et al. Daily Left Prefrontal Transcranial Magnetic Stimulation Therapy for Major Depressive Disorder: A Sham-Controlled Randomized Trial. Archives of General Psychiatry 2010;67:507–16. 10.1001/archgenpsychiatry.2010.46.

[15] Daskalakis ZJ. Repetitive Transcranial Magnetic Stimulation for Major Depressive Disorder: A Review. The Canadian Journal of Psychiatry 2008;53.

[16] Bergmann TO, Lieb A, Zrenner C, Ziemann U. Pulsed Facilitation of Corticospinal Excitability by the Sensorimotor μ-Alpha Rhythm. J Neurosci 2019;39:10034. 10.1523/JNEUROSCI.1730-19.2019.

[17] Madsen KH, Karabanov AN, Krohne LG, Safeldt MG, Tomasevic L, Siebner HR. No trace of phase: Corticomotor excitability is not tuned by phase of pericentral mu-rhythm. Brain Stimulation 2019;12:1261–70. 10.1016/j.brs.2019.05.005.

[18] Karabanov AN, Madsen KH, Krohne LG, Siebner HR. Does pericentral mu-rhythm “power” corticomotor excitability? – A matter of EEG perspective. Brain Stimulation 2021;14:713–22. 10.1016/j.brs.2021.03.017.

[19] Zrenner C, Desideri D, Belardinelli P, Ziemann U. Real-time EEG-defined excitability states determine efficacy of TMS-induced plasticity in human motor cortex. Brain Stimulation 2018;11:374–89. 10.1016/j.brs.2017.11.016.

[20] Schaworonkow N, Caldana Gordon P, Belardinelli P, Ziemann U, Bergmann TO, Zrenner C. μ-Rhythm Extracted With Personalized EEG Filters Correlates With Corticospinal Excitability in Real-Time Phase-Triggered EEG-TMS. Frontiers in Neuroscience 2018;12. 10.3389/fnins.2018.00954.

[21] Schaworonkow N, Triesch J, Ziemann U, Zrenner C. EEG-triggered TMS reveals stronger brain state-dependent modulation of motor evoked potentials at weaker stimulation intensities. Brain Stimulation 2019;12:110–8. 10.1016/j.brs.2018.09.009.

[22] Desideri D, Zrenner C, Ziemann U, Belardinelli P. Phase of sensorimotor μ-oscillation modulates cortical responses to transcranial magnetic stimulation of the human motor cortex. The Journal of Physiology 2019;597:5671–86. 10.1113/JP278638.

[23] Esser SK, Huber R, Massimini M, Peterson MJ, Ferrarelli F, Tononi G. A direct demonstration of cortical LTP in humans: A combined TMS/EEG study. Brain Research Bulletin 2006;69:86–94. 10.1016/j.brainresbull.2005.11.003.

[24] Litvak V, Komssi S, Scherg M, Hoechstetter K, Classen J, Zaaroor M, et al. Artifact correction and source analysis of early electroencephalographic responses evoked by transcranial magnetic stimulation over primary motor cortex. NeuroImage 2007;37:56–70. 10.1016/j.neuroimage.2007.05.015.

[25] Mäki H, Ilmoniemi RJ. The relationship between peripheral and early cortical activation induced by transcranial magnetic stimulation. Neuroscience Letters 2010;478:24–8. 10.1016/j.neulet.2010.04.059.

[26] Petrichella S, Johnson N, He B. The influence of corticospinal activity on TMS-evoked activity and connectivity in healthy subjects: A TMS-EEG study. PLOS ONE 2017;12:e0174879. 10.1371/journal.pone.0174879.

[27] Roos D, Biermann L, Jarczok TA, Bender S. Local Differences in Cortical Excitability – A Systematic Mapping Study of the TMS-Evoked N100 Component. Frontiers in Neuroscience 2021;15. 10.3389/fnins.2021.623692.

[28] Perera ND, Alekseichuk I, Shirinpour S, Wischnewski M, Linn G, Masiello K, et al. Dissociation of Centrally and Peripherally Induced Transcranial Magnetic Stimulation Effects in Nonhuman Primates. J Neurosci 2023;43:8649. 10.1523/JNEUROSCI.1016-23.2023.

[29] Wischnewski M, Haigh ZJ, Shirinpour S, Alekseichuk I, Opitz A. The phase of sensorimotor mu and beta oscillations has the opposite effect on corticospinal excitability. Brain Stimulation 2022;15:1093–100. 10.1016/j.brs.2022.08.005.

[30] P. Julkunen. Mobile Application for Adaptive Threshold Hunting in Transcranial Magnetic Stimulation. IEEE Transactions on Neural Systems and Rehabilitation Engineering 2019;27:1504–10. 10.1109/TNSRE.2019.2925904.

[31] Shirinpour S, Alekseichuk I, Mantell K, Opitz A. Experimental evaluation of methods for real-time EEG phase-specific transcranial magnetic stimulation. J Neural Eng 2020;17:046002. 10.1088/1741-2552/ab9dba.

[32] Oostenveld R, Fries P, Maris E, Schoffelen J-M. FieldTrip: Open Source Software for Advanced Analysis of MEG, EEG, and Invasive Electrophysiological Data. Computational Intelligence and Neuroscience 2011;2011:1–9. 10.1155/2011/156869.

[33] Bell AJ, Sejnowski TJ. An information-maximization approach to blind separation and blind deconvolution. Neural Comput 1995;7:1129–59. 10.1162/neco.1995.7.6.1129.

[34] Ilmoniemi RJ, Kičić D. Methodology for Combined TMS and EEG. Brain Topogr 2010;22:233–48. 10.1007/s10548-009-0123-4.

[35] Kruglikov SY, Schiff SJ. Interplay of Electroencephalogram Phase and Auditory-Evoked Neural Activity. J Neurosci 2003;23:10122–7. 10.1523/JNEUROSCI.23-31-10122.2003.

[36] Bergmann TO, Mölle M, Schmidt MA, Lindner C, Marshall L, Born J, et al. EEG-Guided Transcranial Magnetic Stimulation Reveals Rapid Shifts in Motor Cortical Excitability during the Human Sleep Slow Oscillation. Journal of Neuroscience 2012;32:243–53. 10.1523/JNEUROSCI.4792-11.2012.

[37] Mazaheri A, Jensen O. Posterior α activity is not phase-reset by visual stimuli. Proceedings of the National Academy of Sciences 2006;103:2948–52. 10.1073/pnas.0505785103.

[38] Farzan F, Barr MS, Hoppenbrouwers SS, Fitzgerald PB, Chen R, Pascual-Leone A, et al. The EEG correlates of the TMS-induced EMG silent period in humans. NeuroImage 2013;83:120–34. 10.1016/j.neuroimage.2013.06.059.

[39] Nikulin VV, Kicic D, Kahkonen S, Ilmoniemi RJ. Modulation of electroencephalographic responses to transcranial magnetic stimulation: evidence for changes in cortical excitability related to movement. Eur J Neurosci 2003;18:1206–12. 10.1046/j.1460-9568.2003.02858.x.

[40] Rogasch NC, Fitzgerald PB. Assessing cortical network properties using TMS-EEG. Hum Brain Mapp 2013;34:1652–69. 10.1002/hbm.22016.

[41] Rogasch NC, Daskalakis ZJ, Fitzgerald PB. Mechanisms underlying long-interval cortical inhibition in the human motor cortex: a TMS-EEG study. Journal of Neurophysiology 2013;109:89–98. 10.1152/jn.00762.2012.

[42] Premoli I, Castellanos N, Rivolta D, Belardinelli P, Bajo R, Zipser C, et al. TMS-EEG Signatures of GABAergic Neurotransmission in the Human Cortex. Journal of Neuroscience 2014;34:5603–12. 10.1523/JNEUROSCI.5089-13.2014.

[43] Bonnard M, Spieser L, Meziane HB, De Graaf JB, Pailhous J. Prior intention can locally tune inhibitory processes in the primary motor cortex: direct evidence from combined TMS-EEG. European Journal of Neuroscience 2009;30:913–23. 10.1111/j.1460-9568.2009.06864.x.

[44] Rogasch NC, Biabani M, Mutanen TP. Designing and comparing cleaning pipelines for TMS-EEG data: A theoretical overview and practical example. Journal of Neuroscience Methods 2022;371:109494. 10.1016/j.jneumeth.2022.109494.

[45] Bertazzoli G, Esposito R, Mutanen TP, Ferrari C, Ilmoniemi RJ, Miniussi C, et al. The impact of artifact removal approaches on TMS–EEG signal. NeuroImage 2021;239:118272. 10.1016/j.neuroimage.2021.118272.

[46] Zazio A, Miniussi C, Bortoletto M. Alpha-band cortico-cortical phase synchronization is associated with effective connectivity in the motor network. Clinical Neurophysiology 2021;132:2473–80. 10.1016/j.clinph.2021.06.025.

[47] Weise K, Numssen O, Thielscher A, Hartwigsen G, Knösche TR. A novel approach to localize cortical TMS effects. NeuroImage 2020;209. 10.1016/j.neuroimage.2019.116486.

[48] Salvador R, Silva S, Basser PJ, Miranda PC. Determining which mechanisms lead to activation in the motor cortex: A modeling study of transcranial magnetic stimulation using realistic stimulus waveforms and sulcal geometry. Clinical Neurophysiology 2011;122:748–58. 10.1016/j.clinph.2010.09.022.

[49] Fecchio M, Pigorini A, Comanducci A, Sarasso S, Casarotto S, Premoli I, et al. The spectral features of EEG responses to transcranial magnetic stimulation of the primary motor cortex depend on the amplitude of the motor evoked potentials. PLOS ONE 2017;12:e0184910. 10.1371/journal.pone.0184910.

[50] Peters JC, Reithler J, Graaf TA de, Schuhmann T, Goebel R, Sack AT. Concurrent human TMS-EEG-fMRI enables monitoring of oscillatory brain state-dependent gating of cortico-subcortical network activity. Communications Biology 2020;3:40. 10.1038/s42003-020-0764-0.

[51] Ozdemir RA, Kirkman S, Magnuson JR, Fried PJ, Pascual-Leone A, Shafi MM. Phase matters when there is power: Phasic modulation of corticospinal excitability occurs at high amplitude sensorimotor mu-oscillations. Neuroimage: Reports 2022;2:100132. 10.1016/j.ynirp.2022.100132.

[52] Kilavik BE, Zaepffel M, Brovelli A, MacKay WA, Riehle A. The ups and downs of beta oscillations in sensorimotor cortex. Experimental Neurology 2013;245:15–26. 10.1016/j.expneurol.2012.09.014.

[53] Jana S, Hannah R, Muralidharan V, Aron AR. Temporal cascade of frontal, motor and muscle processes underlying human action-stopping. eLife 2020;9:e50371. 10.7554/eLife.50371.

[54] Swann NC, Cai W, Conner CR, Pieters TA, Claffey MP, George JS, et al. Roles for the pre-supplementary motor area and the right inferior frontal gyrus in stopping action: Electrophysiological responses and functional and structural connectivity. NeuroImage 2012;59:2860–70. 10.1016/j.neuroimage.2011.09.049.

[55] Wagner J, Wessel JR, Ghahremani A, Aron AR. Establishing a Right Frontal Beta Signature for Stopping Action in Scalp EEG: Implications for Testing Inhibitory Control in Other Task Contexts. Journal of Cognitive Neuroscience 2018;30:107–18. 10.1162/jocn_a_01183.

[56] Kawasaki M, Uno Y, Mori J, Kobata K, Kitajo K. Transcranial magnetic stimulation-induced global propagation of transient phase resetting associated with directional information flow. Frontiers in Human Neuroscience 2014;8. 10.3389/fnhum.2014.00173.

[57] Paus T, Sipila PK, Strafella AP. Synchronization of Neuronal Activity in the Human Primary Motor Cortex by Transcranial Magnetic Stimulation: An EEG Study. Journal of Neurophysiology 2001;86:1983–90. 10.1152/jn.2001.86.4.1983.

[58] Hanslmayr S, Klimesch W, Sauseng P, Gruber W, Doppelmayr M, Freunberger R, et al. Alpha Phase Reset Contributes to the Generation of ERPs. Cerebral Cortex 2007;17:1–8. 10.1093/cercor/bhj129.

[59] Gruber WR, Klimesch W, Sauseng P, Doppelmayr M. Alpha Phase Synchronization Predicts P1 and N1 Latency and Amplitude Size. Cerebral Cortex 2005;15:371–7. 10.1093/cercor/bhh139.

[60] Di Lazzaro V, Oliviero A, Profice P, Saturno E, Pilato F, Insola A, et al. Comparison of descending volleys evoked by transcranial magnetic and electric stimulation in conscious humans. Electroencephalogr Clin Neurophysiol 1998;109:397–401. 10.1016/s0924-980x(98)00038-1.

