## Supplemental Data for "State dependent motor cortex stimulation reveals distinct mechanisms for corticospinal excitability and cortical responses"

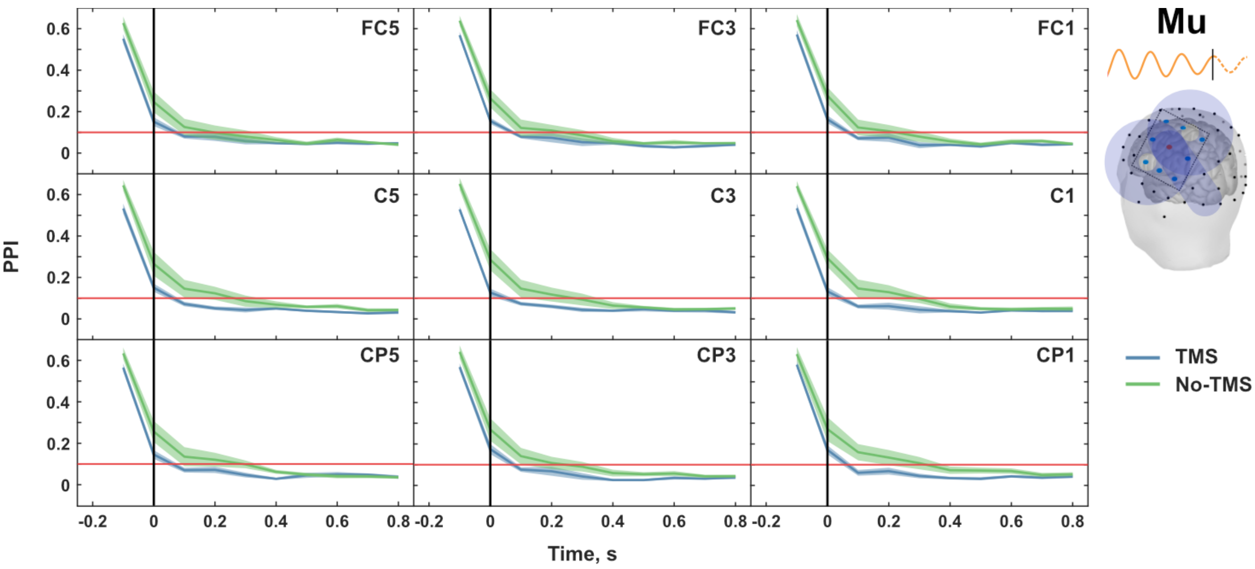


***Figure 1. Phase Preservation Index (PPI) for mu-specific targeting.*** *PPI values are calculated with reference to mu phase at 100 ms prior to TMS pulse, at 100 ms intervals. Blue trace indicates the PPI values for real TMS, and green trace indicates PPI for TMS trigger without actual pulse. Shaded region depicts the standard error of mean (SEM) of PPI. Plots for all electrodes in the Laplacian montage (FC5, FC3, FC1, C5, C3, C1, CP5, CP3 and CP1) are shown here. Black vertical line indicates the TMS trigger, and the red horizontal line (PPI = 0.1) indicates the threshold for phase preservation calculated according to Fischer, 1993. For real TMS condition, there is no substantial evidence for mu phase preservation after TMS delivery.*

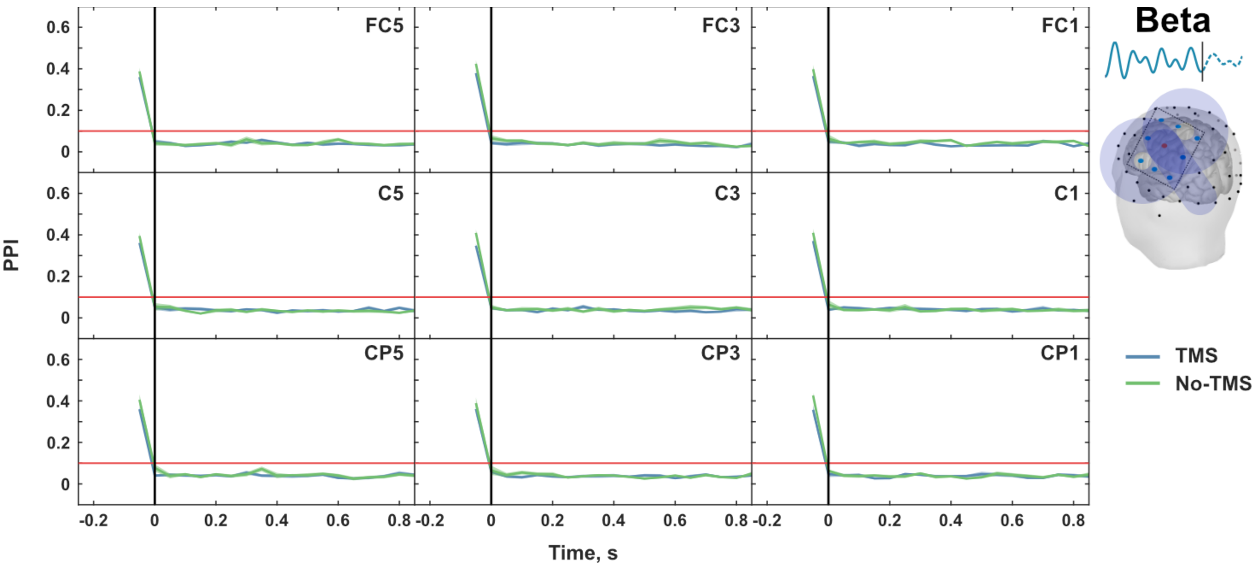


***Figure 2. Phase Preservation Index (PPI) for beta-specific targeting.*** *PPI values are calculated with reference to beta phase at 100 ms prior to TMS pulse, at 100 ms intervals. Blue trace indicates the PPI values for real TMS, and green trace indicates PPI for TMS trigger without actual pulse. Shaded region depicts the standard error of mean (SEM) of PPI. Plots for all electrodes in the Laplacian montage (FC5, FC3, FC1, C5, C3, C1, CP5, CP3 and CP1) are shown here. Black vertical line indicates the TMS trigger, and the red horizontal line (PPI = 0.1) indicates the threshold for phase preservation calculated according to Fischer, 1993. For real TMS condition, there is no substantial evidence for beta phase preservation after TMS delivery.*
